# A thermodynamic model for interpreting tryptophan excitation-energy-dependent fluorescence spectra provides insight into protein conformational sampling and stability

**DOI:** 10.1101/2021.09.09.459605

**Authors:** A Kwok, IS Camacho, S Winter, M Knight, RM Meade, MW Van der Kamp, A Turner, J O’Hara, JM Mason, AR Jones, VL Arcus, CR Pudney

## Abstract

It is now over thirty years since Demchenko and Ladokhin first posited the potential of the tryptophan red edge excitation shift (REES) effect to capture information on protein molecular dynamics. Whilst there have been many key efforts in the intervening years, a biophysical thermodynamic model to quantify the relationship between the REES effect and protein flexibility has been lacking. Without such a model the full potential of the REES effect cannot be realized. Here, we present a thermodynamic model of the protein REES effect that captures information on protein conformational flexibility, even with proteins containing multiple tryptophan residues. Our study incorporates exemplars at every scale, from tryptophan in solution, single tryptophan peptides to multi-tryptophan proteins, with examples including a structurally disordered peptide, *de novo* designed enzyme, human regulatory protein, therapeutic monoclonal antibody in active commercial development, and a mesophilic and hyperthermophilic enzyme. Combined, our model and data suggest a route forward for the experimental measurement of the protein REES effect and point to the potential for integrating bimolecular simulation with experimental data to yield novel insights.

---

Tracking protein conformational change, and even more subtly, changes in the equilibrium of available conformational states is central to molecular biosciences. Protein stability is intimately linked with the distribution of conformational states^1^ and as a good generalisation, increased stability tracks with a decrease in the distribution of conformational states (increasing rigidity).^2^ While engineering protein stability has advanced enormously, the tools to sensitively and quantitatively track these changes are lacking. There are a broad range of potential analytical tools, but only a few which can be applied routinely to the vast majority of proteins without unreasonable requirements regarding solvent, protein concentrations and thermal stability, or without the requirement of surface attachment or labelling.^3^ Moreover, the vast majority of protein conformational changes are subtle, described as ‘breathing’ motions, where most structural orders (primary to quaternary) of the protein are not altered, but it is the equilibrium of conformational states (protein flexibility) that changes.^4^

The red edge excitation shift (REES) phenomenon is a sensitive reporter of a fluorophore’s environment and the mechanism is shown in Figure 1A.^5–8^ Radiative fluorescence takes place after light absorption alongside two non-radiative processes, which include vibrational relaxation and solvent relaxation (dipolar re-organisation). Vibrational relaxation is typically fast (~10^−12^ s) relative to the lifetime of fluorescence emission (*τ*F ~ 10^−10^ - 10^−9^ s) and so causes a complete relaxation of the system to its lowest energy level prior to emission. This gives rise to the familiar red shift of a fluorescence emission compared to absorption (Stokes shift). The Lippert-Mataga equation illustrates that the greater the polarity of solvent, the larger the anticipated Stokes shift.^9–10^

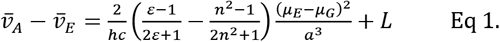

where the Stokes shift (wavenumber of absorption and emission), 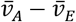, is governed by the dielectric of the solvent, *∊*, specifically the reorientation of solvent dipoles; the refractive index, *n;* the dipole moment of the ground and excited states, *μ_G_* and *μ_E_*, respectively; the radius of the fluorophore cavity, *a*; and a constant, *L.*

**Figure 1.**
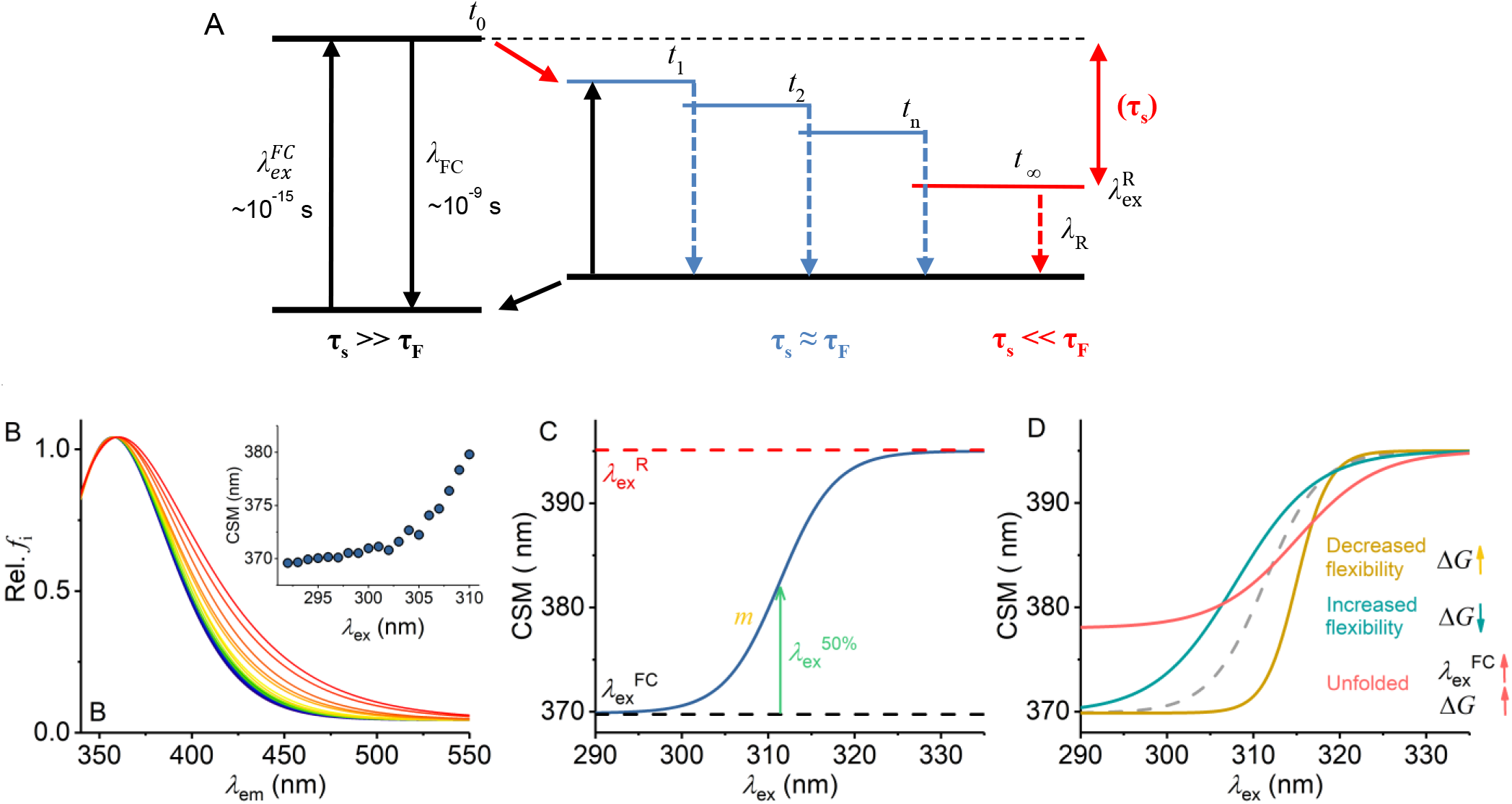
Mechanism of the REES effect, predicted experimental observables and graphical description of Eq 7. **A,** Jablonski diagram illustrating the REES effect. **B,** Example model Trp REES data showing the normalized emission spectrum with increasing excitation wavelength and *inset* as the change in CSM *versus* excitation wavelength. **C,** Graphical depiction of Eq 7. **D,** Predicted spectral changes resulting from variations in Eq 7 from shifts in protein flexibility and conformational state (folding).

Eq 1 assumes that the solvent relaxation is complete prior to emission. However, solvent relaxation is not necessarily always fast relative to fluorescence emission and under a range of solvent or environmental conditions can approach the lifetime of fluorescence emission (~10-^10^ - 10^−9^ s). The longer *τ*S can therefore affect the level from which emission occurs and so the emission wavelength, in which case it also contributes to the Stokes shift.^5–8^ Specifically, one anticipates an ensemble of energetic sub-states are formed related to the distribution of solvent relaxation lifetimes; i.e., the available distribution of solvent-fluorophore interaction energies. The additive contribution of these states to the steady-state emission spectrum gives rise to broadband emission, which is observed as inhomogeneous broadening of the spectra. This broadening is then dependent on the excitation energy used, since as one decreases the excitation energy there is an increasing photoselection of states (Figure 1A). Experimentally, one then observes a red shift in the emission spectra with respect to increasing excitation wavelength, i.e., decreasing excitation energy (Figure 1B). The inhomogeneous broadening will be dependent on a range of physical conditions that affect *τ*S, including temperature, viscosity and solvent dipole moment (and therefore solvent dielectric). ^5–8^

The sensitivity of the REES effect to changes in the equilibrium of solvent-fluorophore interaction energies suggests potential in using the approach to track changes in protein conformational state using the intrinsic fluorescence of the aromatic amino acids.^8,11^ Indeed, tryptophan (Trp) has been shown to give a large REES effect in numerous proteins and we point to the excellent review by Chattopadhyay (ref 11), which illustrates key examples. Demchenko and Ladokhin^12^ suggest that the selection between ^1^L_a_ and ^1^L_b_ electronic excited states acts to increase the magnitude of the red edge excitation shift. Trp has the advantage that it’s emission can be separated from tyrosine (Tyr) and phenylalanine (Phe) by excitation at wavelengths > ~292 nm.^13^ Trp REES is therefore a potentially excellent probe of protein conformational change, and possibly even of changes in the equilibrium of conformational states.

We have previously applied and validated an empirical model for describing protein REES data as a function of the equilibrium of conformational states, which we call QUBES (quantitative understanding of biomolecular edge shift).^14–16^ Herein, we refer to changes in the equilibrium of conformational states as changes in flexibility, with a more flexible protein having a broader equilibrium of conformational states. We track the changes in inhomogeneous broadening as the change in the centre of spectral mass (CSM) of steady-state emission spectra (example shown in Figure 1B),

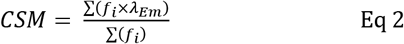

The resulting data are then fit to the QUBES model.

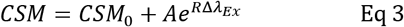

where *CSM*_0_ is the CSM value independent of *λ*_*Em*_. The am-plitude relative to CSM_0_ and curvature of the exponential is described by *A* and *R,* respectively. We have previously found the parameters from this empirical model could be used to track changes in protein stability.^14,16,17^ That this simple model appears to provide useful insight suggests it is approximating the protein REES effect to a level of accuracy.

Whilst Eq 3 performs well at tracking shifts in protein rigidity/flexibility (also for multi-Trp containing proteins),^14–17^ it does not relate to a specific thermodynamic parameter and neglects the fact that protein Trp emission will have a finite maximum observable spectral red shift at 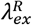. Moreover, the data from our QUBES model cannot be cross compared to proteins with different Trp content and location in structure. Developing our QUBES model towards an accurate *a priori* thermodynamic model would therefore enhance both the accuracy and utility of the approach for studying protein dynamics/stability.

Herein, we describe a thermodynamic model for interpreting protein REES data, which builds on our early work. Using a range of model systems from Trp/solvent studies, single Trp containing proteins and multi-Trp proteins, we find that the new model accurately tracks with independent metrics of changes in the equilibrium of protein conformational states as well as more gross metrics of protein folding. Moreover, our model points to the need for new experimental approaches to monitor the protein REES effect.

## Results and Discussion

As described by Demchenko and Ladokhin^12^ we posit a two-state model and assume [*FC*] ⇌ [*R*] and *τ_F_* ≪ *τ_S_*, then the fractional concentration of *R* is given by:

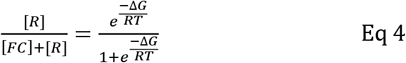

where ΔG is the difference in free energy between the *FC* and *R* states. We then assume that Δ*G* will change linearly with excitation wavelength:

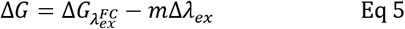

Thus, we anticipate a two-state transition between *FC* and *R* states due to photoselection by excitation wavelength with baselines 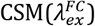 and 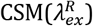. The gradient of the of tran-sition is given by |Δ*G*| at any particular *λ_Ex_*.

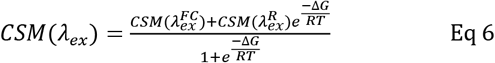

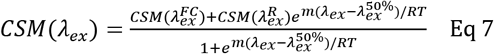

Eq 6 and 7 establish three key parameters 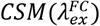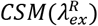, and Δ*G*, which we describe below. Figure 1C shows Eq 7 plotted in a similar manner to the experimental data as in Figure 1B *inset* but now showing the full range of the function (Eq 7). Eq 7 is a more complete description of the REES effect (*c.f*. Eq 3) since it predicts a maximum magnitude of the CSM, corresponding to the fully relaxed state, 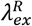 (Figure 1C). Indeed, ourselves and others have observed saturation of the REES effect with non-Trp fluorophores used as molecular probes^18^ or ligands,^19^ and so Eq 7 is logical for the REES effect in proteins. 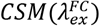 is the CSM corresponding to 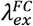 shown in Figure 1A. We anticipate 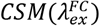 will be responsive to changes in solvation environment in a similar way to the spectral shift of Trp on solvent/exposure/occlusion. That is, as the Burstein classification^20^ and Eq 1, increasing solvent exposure will cause 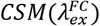 to red-shift and a decrease in solvent exposure will cause 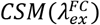 to blue shift.^20^

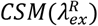 is the CSM corresponding to 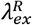 in Figure 1A; *i.e*., the completely relaxed state of the solvent. Note that this value should be fixed for a given system, unlike 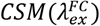, which will be responsive to variation in the solvent environment. This parameter therefore represents entirely novel information over previous models of the REES effect. Spe-cifically, 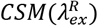 reports on an extreme of the solvent-fluorophore interaction energy. It can therefore be considered a unique identifying parameter related to both protein structure and physiochemical environment.

The combination of 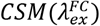 and 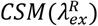 will therefore be a unique measurement of the accessible equilibrium of protein conformational states and will be specific to a specific protein structure, molecular flexibility and Trp content and location.

Δ*G* arises from Eq 4, calculated from the extracted 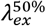 and *m* terms in Eq 6. Where 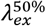 is the *λ_Ex_* at half the maximal CSM and *m* reflects information on the slope of the plot shown in Figure 1C. This gives Δ*G* (J mol^−1^) at a specific wavelength, which has a linear relationship to *λ_Ex_* (Eq 5). For consistency, we report the gradient of this plot of Δ*G versus λ_Ex_*, giving Δ*G* expressed in J mol^−1^ nm^−1^. Δ*G* reports on the energy gap between adjacent emissive states. For example, in the most extreme case, the gap between the *FC* and *R* states as shown in Figure 1A. As the number of intermediate state increases, reflecting an increased distribution of solvent-fluorophore interaction energies, so the magnitude of Δ*G* will increase, representing a broader distribution of intermediate states.

Inspection of Figure 1A yields two ready predictions for the information content of the parameters in Eq 7 and we show how these are predicted to affect the resulting experimental data in Figure 1D:

i. A decrease in the gap between 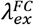 and 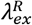 (*i.e.*, an increase in 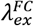) would reflect a narrower distribution – but unchanged number – of solvent-fluorophore interaction energies. That is, based on Hammond’s postulate,^21^ the environments of the *FC* and *R* states becoming more similar. Experimentally this would manifest as an increase in the extracted magnitude of 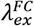 since 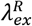 will be a fixed value for a given solvent-fluorophore environment.
ii. In a more rigid molecule we expect to observe fewer intermediate states. Fewer energetically discrete solvent-fluorophore environments would reflect a larger energy gap between adjacent states (*t*_1_, *t*_2_ etc, Figure 1A) and a smaller distribution of solvent-solute interaction energies and would manifest as reduced inhomogenous broadening of the emission spectra (Figure 1C). Experimentally, one then expects a steeper transition between 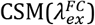 and 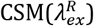, giving rise to an increased Δ*G*.

Changes in both 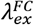 and Δ*G* are possible and indeed likely when studying proteins. As a specific case, for a completely unfolded *versus* folded protein, we anticipate an increase in 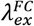 and an increase in Δ*G*. That is, 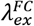 increases due to the increase in solvent exposure of the available Trp residues and Δ*G* increases as the number of intermediate (discrete) solvent-fluorophore interaction energies decrease, tending towards the homogenous single state where all Trps are completely solvent exposed, i.e. as in (i) where the environments of the *FC* and *R* states become more similar.

We acknowledge that it is not possible to experimentally reach saturation of the Trp REES effect 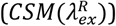 using conventional spectrometers owing to the technical limitations of the intensity of UV light (using halogen lamps) as well as convolution of the emission spectra with the relatively broad-band excitation achieved from monochromation at the large slit widths necessary to increase illumination. In practice we find the signal to noise ratio becomes intractable beyond *λ_ex_* ≈ 310 nm for the same concentration of protein. We discuss this in more detail below.

### Tryptophan in solution

Given that Eq 7 is a new thermodynamic model for the REES effect we first explore the sensitivity of the Trp REES effect to variation in the physical properties of the solvent. Solvent studies have been used to probe the sensitivity of the REES effect using viscous matrices such as ethylene glycol and glycerol as well as temperature variation, by monitoring Trp or indole emission.^7,12^ One expects the REES effect to be sensitive to changes in the dielectric and viscosity of the solvent and the temperature owing to the effect on the lifetime of solvent relaxation as described above. We are not aware of a method to independently vary dielectric, viscosity and temperature so we have employed a matrix effect experiment, where we monitor the Trp REES effect as a function of methanol (MeOH) concentration (0-70% v/v) and temperature (20-50 °C). Figure S1 shows the variation in viscosity and dielectric for the conditions we use. Using this approach we are able to explore the REES effect, which is quantified using Eq 6 across a this range of conditions. Figures 2A-D show the raw REES data as a function of the variation in MeOH concentration at each temperature studied. These data are then fit to Eq 7 and the resulting parameters are shown in Figures 2E-G.

**Figure 2.**
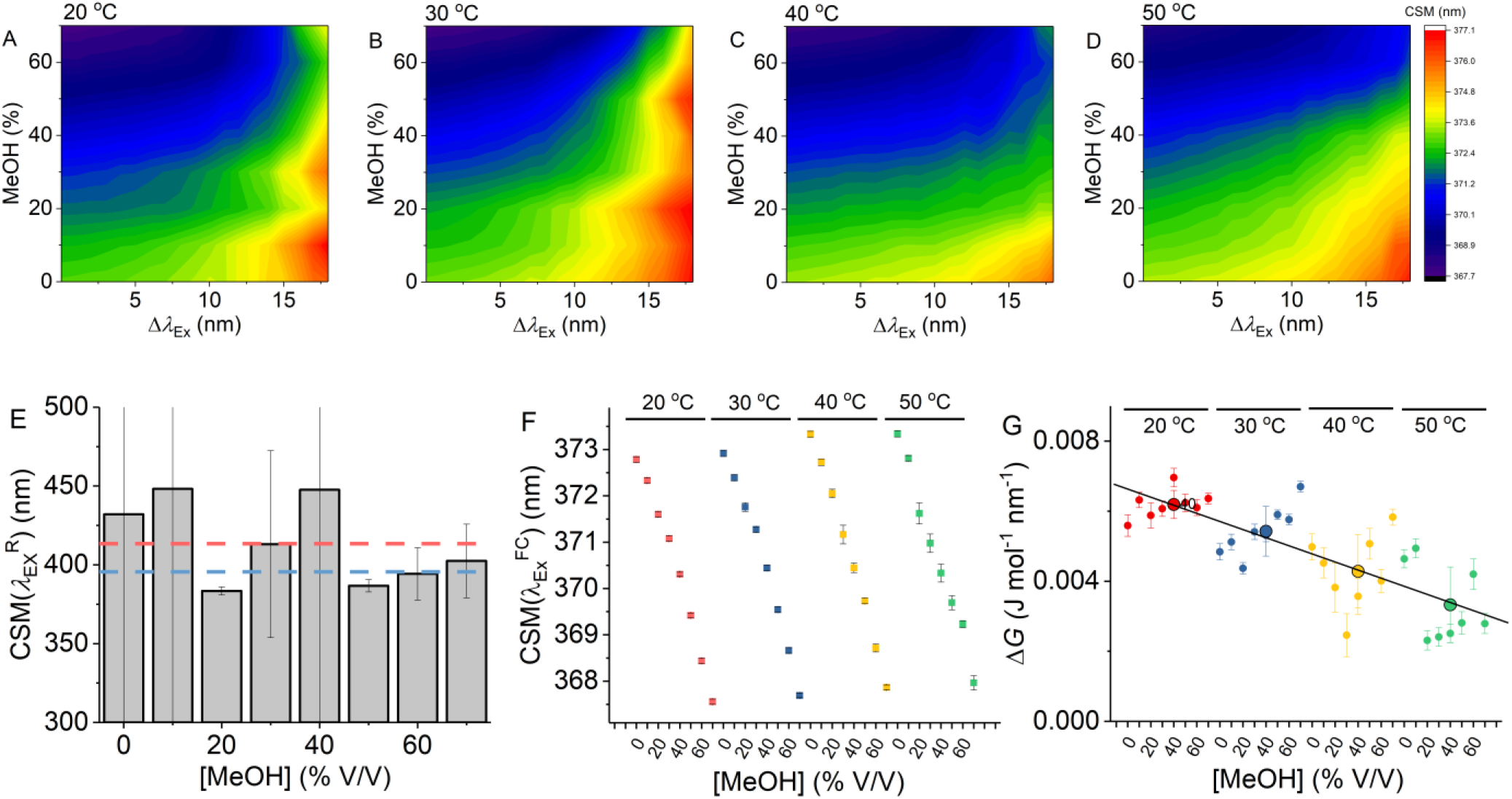
Solvent and temperature studies of the Trp REES effect. **A-D,** Variation in CSM for L-Trp with varying percentages of MeOH and *versus* temperature. **E,** Variation of the 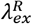 value for each [MeOH] studied, where the fitted 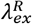 value is a shared parameter for each temperature at a given [MeOH]. The dashed red line shows the average 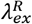 value and the blue dashed line shows the 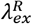 value extracted where all data are fit to Eq 7 with 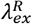 as a shared parameter. **F,** Variation of the 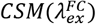 value for each condition studied. **G,** Variation of the Δ*G* at fixed 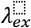 value for each condition studied. Large coloured dots represent the average of [MeOH] at each temperature and the error bars are the standard deviation. *Conditions*, 1 μM L-Trp, 50mM Tris-HCL, pH 8.0.

As we describe above, accessing the limiting value of 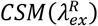 experimentally is challenging and thus the extracted value of 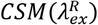 from fits to Eq 7 will necessarily have a large error and in some cases, the extracted values are unrealistically large (> 1000 nm). As an alternative one can share the value of 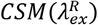 during fitting, which provides much greater restraint as well as improved accuracy on the extracted magnitude of 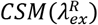. Fitting with 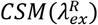 as a shared parameter for all the data sets gives an average and standard deviation of 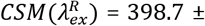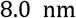 (Figure 2E). However, we are aware this likely masks much of the real variation in the magnitude of 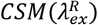, not least because we expect variation in this parameter with changes in dielectric. Alternatively, fitting the data with shared values of 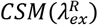 for the same [MeOH] but at varying temperatures (Figure 2E) gives 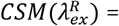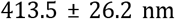. These data suggest a practical range of 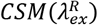 (at least across the range of the conditions explored in Figure 2) from ~387 to ~440 nm.

Figure S2 shows modelled data showing the effect of varying 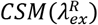 on the extracted magnitude of Δ*G* (there is no effect on 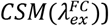). These data show a ~ 10% variance in Δ*G* across this range of 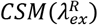 values tested and so the effect of using a fixed value of 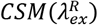 is not large. We note hat the range of dielectric and viscosity values this represents is far broader than for a protein in aqueous solvent. Therefore, whilst not ideal, until it is experimentally possible to extract data at very low excitation energies (>*λ_ex_* = 310 nm), fixing the magnitude of 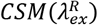 is necessary to extract realistic values for Δ*G* and our data imply this will not cause a large effect on protein data. We therefore use 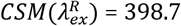 (as above) to extract values of 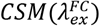 and Δ*G* for the data shown in Figure 2F and 2G.

Figure 2F shows the variation in 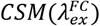 for each [MeOH] at each temperature studied. At all temperatures the magnitude of 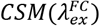 decrease with increasing [MeOH]. This decrease is expected for a simple solvatochromatic shift and has been observed in numerous cases previously. This expected finding is satisfying because it validates the interpretation of 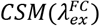 value as an excitation wavelength-independent metric of Trp solvation. Figure S3 shows the temperature dependence of 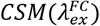 at each [MeOH], extracted from fitting to a simple linear function. Figure S3 shows a ‘V-shaped’ temperature dependence with respect to [MeOH], with a minimum at 30% [MeOH], where there is no measurable temperature dependence of 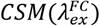 within error. Therefore, our data suggest that in aqueous solvent, 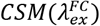 appears to have an intrinsic temperature dependence of ~ 0.02 nm^−1^ K^−1^ for free Trp in aqueous solution. We consider whether this is borne out in protein samples below.

Figure 2G shows the variation in the extracted magnitude of Δ*G* as a function of [MeOH] at each temperature studied. We find a general decrease in the magnitude of Δ*G* with increasing temperature (−0.1 × 10^−3^ J mol^−1^ nm-^1^ K^−1^ across the range studied). Increased temperature will increase τS thus, one anticipates a smaller REES effect and, as described above, a decrease in the magnitude of Δ*G* as we indeed observe. That our data track with a logical and expected physical effect validates the principles used to derive Eq 7.

From Figure 2G we do not observe a consistent trend in the magnitude of Δ*G* with respect to [MeOH]. It is not possible to independently vary viscosity, dielectric and temperature, with viscosity having a strong dependence on both temperature and [MeOH]. In contrast to 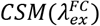, it is evident that Δ*G* is acutely sensitive to such inter-dependencies. it is therefore not possible to assess simple trends in Δ*G* as a function of [MeOH]. To illustrate this point, we have plotted the magnitude of Δ*G versus* the calculated solvent viscosity and dielectric for the combination of [MeOH] and temperature used – Figure S1C. From this figure, it is apparent that there is a complex trend governing the magnitude of Δ*G*, resembling an elliptical phase-type relationship. What these data do serve to illustrate, is not only the extreme sensitivity of the REES effect to the solvent environment as predicted, but also the potential sensitivity of Eq 7 to track these subtle changes in the distribution of solvent’ solute interaction energies.

Our data using free Trp in solution provides a detailed baseline for the sensitivity of Eq 7 to track the protein Trp REES effect, most notably establishing realistic ranges for the magnitude of CSM(λ_ex^R), the temperature dependence of CSM(λ_ex^FC) and illustrating the extreme sensitivity of the magnitude of ΔG to a change in the solvent-fluorophore interaction energies. We wished to directly validate the saturation of CSM (CSM(λ_ex^R)) as shown in Figure 1C and to confirm that the extracted value of CSM(λ_ex^R) = 398.7 from Figure 2 is an accurate reflection of CSM(λ_ex^R). As we discuss above, there are significant technical challenges in collect a ‘complete’ REES data set (measuring emission spectra at λEx > 310 nm). However, using a combination of elevated L-Trp concentration (1.25 mM), a non-aqueous solvent (100% MeOH) and high excitation power (~100 μW), we have achieved this goal, shown in Figure 3. Figure 3A shows the averaged raw spectral data. CSM is calculated in the range 340-500 nm to be consistent across all excitation wavelengths used without being convolved of excitation peaks. From Figure 3B, the resulting CSM data saturate as predicted by Eq 6 and fitting using Eq 7 gives CSM(λ_ex^R) = 397.8 ± 4.0. This compares with CSM(λ_ex^R) = 398.7 ± 8.0 nm extracted from fitting to the Trp REES data (Figure 2) as described above. That these values are effectively identical is powerful validation that the CSM(λ_ex^R) value extracted from simultaneous fitting of REES data (Figure 2) is accurate. To our knowledge this is the first experimental measurement of a complete REES data set. However, we note the conditions used (very high concentration and non-aqueous solvent) are not practical for proteins and we consider alternative routes to achieve this below.

**Figure 3.**
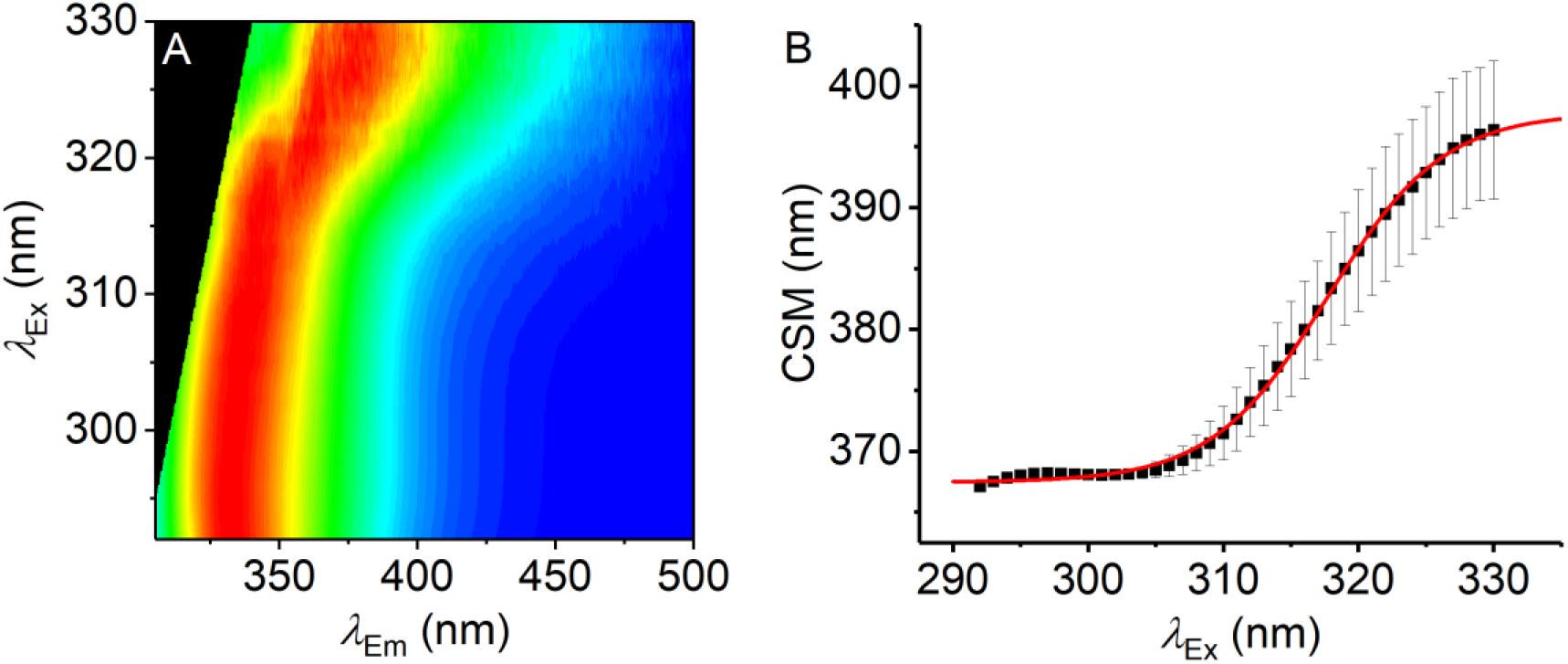
The REES effect monitored over extended λEx range. **A,** Averaged, normalized raw emission spectra. **B,** CSM data extracted from lEm = 340-500 nm. The solid red line is the fit to Eq 7. Data collected in triplicate, error bars are the standard deviation. *Conditions*, 1.25 mM L-Trp, MeOH, 25 °C.

### Single Trp proteins

With the characterization of the REES effect for free Trp in solution in hand, we now turn to single Trp-containing proteins to establish how the REES effect (quantified with Eq 7) changes when the Trp is part of a complex polymer (protein). We have selected a large, monomeric (48 kDa; 419 aa) human regulatory protein, which natively has a single Trp (NF-κB essential modulator - NEMO)^22^, and a natively unstructured protein (α-synuclein, 140 aa)^23^ that lacks native Trp residues, but where we have engineered them into specific sites. These model systems allow us to explore a broad range of conditions and physical environments for single Trp proteins. It also enables us both to explore the sensitivity of Δ*G* but also, similar to our Trp in solution studies, define the range of 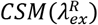 magnitudes for protein/peptides in an aqueous environment *versus* the much broader range of physical conditions studied for Trp in MeOH/water mixtures as described above.

Figure 4A shows a structural model of α-synuclein, with the location of the selected sites for Trp incorporation. α-synuclein is thought to be a largely unstructured (lacking secondary structure) monomer, but which organises into a β-sheet rich fibrillar-like architecture as a repeating unit with a ‘Greek Key’ motif (Figure 4A).^24^ The Trp incorporation sites were selected because in previous work they were found not to alter the aggregation propensity of α-synuclein but did show a measurable REES effect.^25^ In addition, we show data for α-synuclein in the presence and absence of a therapeutic peptide (KDGIVNGVKA), designed to prevent aggregation to the toxic species (as we have reported previously).^26^ This peptide is based on residues 45-54 of the α-synuclein sequence (Figure 4A; green colouration) and therefore binding will be in that location.^26^ This peptide has been shown to bind to a partially aggregated form of α-synuclein.^26^

**Figure 4.**
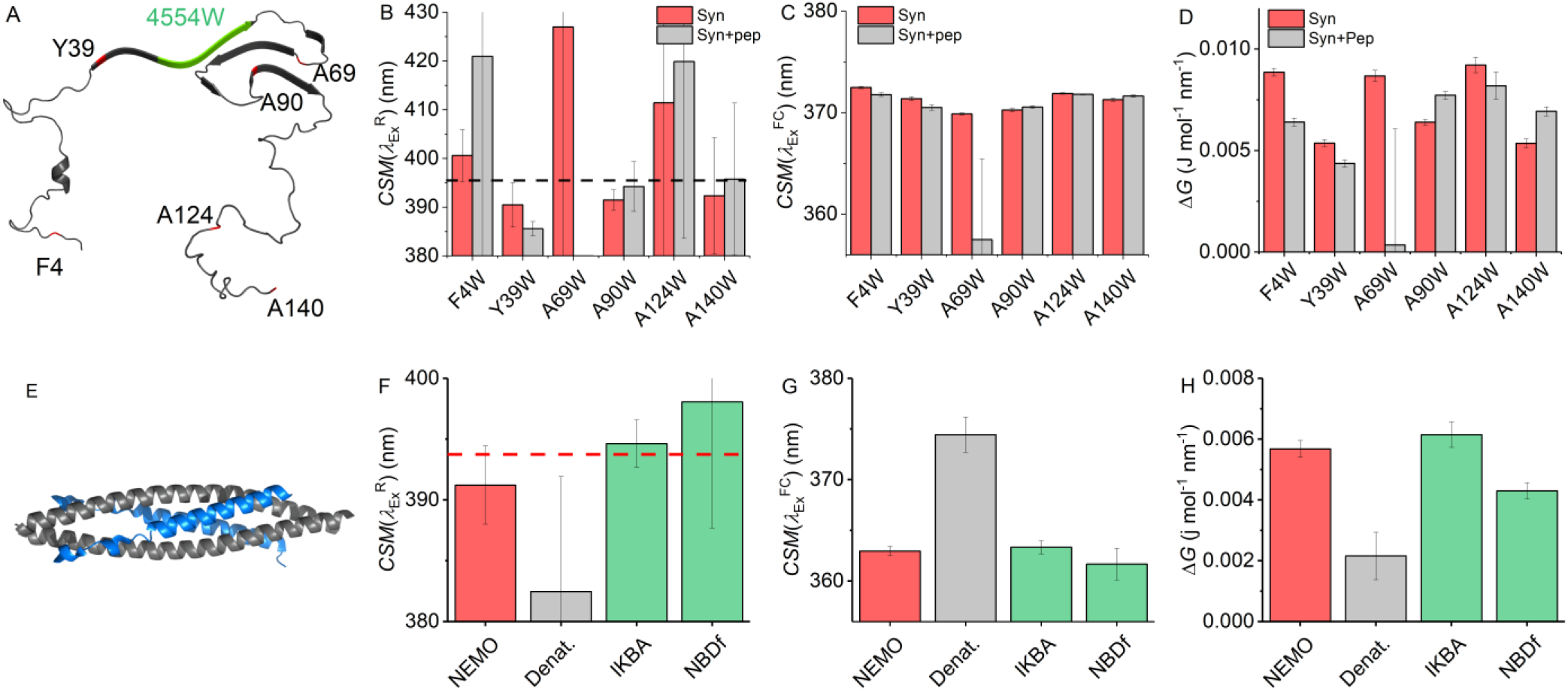
Single Trp protein REES. **A,** Structural model of α-synuclein (PDB 2n0A^24^). **B-C,** Variation in 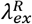 (**B**), 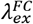 (**C**) and Δ*G* (**D**) extracted from fits of raw REES data to Eq 7 for α-synuclein (red bars) and in the presence of the therapeutic peptide (grey bars). The black dashed line in panel B shows the 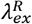 value extracted where all data are fit to Eq 7 with 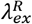 as a shared parameter. **E,** Structural model of the N-terminus of NEMO (3brv) in complex with a peptide representing IKKβ (blue). *Conditions,* 5 μM α-synuclein, 50mM Tris-HCL, pH 8.0. **F-H,** Variation in 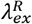 (**B**), 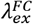 (**C**) and Δ*G* (**D**) extracted from fits of raw REES data to Eq 7 for NEMO (red bars), under denaturing conditions (grey bars) and in the presence of ligands (green bars). The red dashed line in panel F shows the 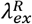 value extracted where all data are fit to Eq 7 with 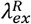 as a shared parameter. Raw NEMO REES data as previously reported in ref 28.

Figure 4B shows the 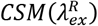 value extracted from the REES data from independent fits (no shared 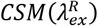 value) to each of the α-synuclein variants and in the presence of the therapeutic peptide. The 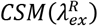 values vary between ~385 and ~425 nm (noting the very large attendant error values in Figure 4B) with an average 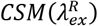 of 400.4 ± 15.4 nm. Sharing the value of 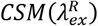 during the fitting to Eq 7, gives 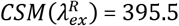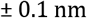. It is worth noting these values of 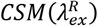 are effectively identical to those extracted for Trp in solution (Figure 2). For consistency in our data analysis we have used 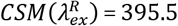 to extract the magnitude of 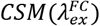 and Δ*G*, as discussed above.

Figure 4C shows the extracted 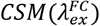 values for each variant and with and without the therapeutic peptide bound. The magnitude of 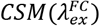 shows variation with Trp position, likely reflecting the combination of the difference in solvent exposure and the immediate electronic environment arising from differences in amino acid composition flanking each Trp. As discussed above, this is effectively a solvatochromatic effect as is typical of Trp emission. However, in the presence of the therapeutic peptide, we find a substantial shift to a lower wavelength for A69W, suggesting a significant decrease in solvent exposure at residue 69 upon peptide binding.

Figure 4D shows the resulting Δ*G* values at each site, extracted from fitting the REES data to Eq 7. We find that the magnitude of Δ*G* varies depending on the specific Trp location in the α-synuclein peptide, which potentially points to some non-globular local structural organisation, similar to a molten globule-like protein. Alternatively, the differences might be attributable to the specific amino acid sequence immediately flanking these positions providing a different distribution of solvent-fluorophore interaction energies. Also, addition of the therapeutic peptide decreases the magnitude of Δ*G* most significantly at a single site, residue 69, similar to our findings for 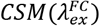.

The finding of a decrease in both Δ*G* and 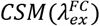 at AA69 on peptide binding suggests that incubation with the therapeutic peptide decreases solvent exposure and increases flexibility at AA69. From Figure 34, AA69W is the variant that is most structurally localised with the anticipated binding site of the therapeutic peptide (green colourisation in Figure 4A). Therefore, our finding of a decreased solvent exposure and shift in flexibility at AA69 is entirely consistent with the putative binding location and the disruption of the putative Greek key motif. These data are powerful evidence that the REES effect, quantified with Eq 7 could be used to track ligand binding and specifically protein-protein interactions.

NF-κB essential modulator (NEMO) is a 48 kDa human regulatory protein involved in the mediation of the NF-κB signalling pathway. A range of studies suggest that NEMO is a flexible protein and can undergo ligand-specific conformational change.^15,27^ It has a single native Trp residue (W6), which is conveniently located close to the residues that bind to the kinase regulated by NEMO (Figure 4E), IκB kinase-β (IKK-β).^22^ Moreover, there is evidence that the IKK-β substrate, IκBα, is also able to interact with NEMO.^28^ We have previously reported the binding of peptide mimics of these proteins to NEMO. We note that the peptides lack Trp residues either natively (IKBα) or by design (NBDPhe, where the native Trp of the NEMO biding domain (NBD) of IKK-β is replaced by Phe).^15^ Figure 4F-H show the results of fitting Eq 7 to NEMO REES data in native and denatured forms, and in the presence of these two ligands.

From Figure 4F we find that the extracted magnitude of 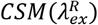 is similar for the different conditions we study (denatured in 8 M urea and with different ligands bound), though we acknowledge that the attendant error is very large (Figure 4E). As with α-synuclein we fit the combined data to Eq 7 but sharing the 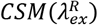 parameter which gives, 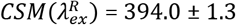. As with α-synuclein, we use this value for 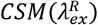 to extract the magnitude of Δ*G* for NEMO.

From Figure 4G we find that the magnitude of 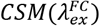 is similar within error for NEMO with and without ligands bound. However, for the unfolded protein in 8M urea, we find that 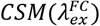 increases from 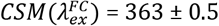 to 374.4 ± 1.7 nm. As we discuss above, the magnitude of 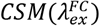 appears to reflect the degree of solvent exposure to the aqueous environment. Therefore, the observation of an increase in 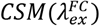 in the presence of denaturant NEMO and with ligands bound. These data show a decrease in Δ*G* when NEMO is denatured (ΔΔ*G* = 0.002 ± 0.001 J mol^−1^ nm^− 1^), no change outside of error in the presence of IKBα (ΔΔ*G* = 0.004 ± 0.0004 J mol^−1^ nm^−1^) and a decrease with NBD-Phe bound (ΔΔ*G* = 0.004 ± 0.0003 J mol^−1^ nm^−1^).

Combined, our data provide a means to interpret the physical meaning of the magnitude of Δ*G*. In the case of the denatured NEMO, the increase in 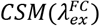 reflects the unfolding of NEMO as an increase in aqueous solvent exposure of the single native Trp residue. The observation of a decrease in the magnitude of Δ*G* would seem consistent with a more heterogeneous (less folded) protein. Binding of NBD-Phe similarly decreases the magnitude of Δ*G* but to a much lesser extent than for unfolded NEMO. Moreover, unlike in the case of the unfolded protein, the magnitude of 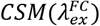 is essentially invariant within error. These data would then suggest a structurally similar protein, but with a partially restricted distribution of conformational states; arguably more ‘folded’ than NEMO alone. This inference seems credible since binding of NEMO to IKKβ gives a well folded α-helical dimer (Figure 3E), despite the binding interface being highly dynamic.^22^ Moreover, these findings track with the binding of the therapeutic peptide to α-synuclein, which shows a similar decrease in the magnitude of Δ*G* on ligand binding (discussed above).

NEMO and α-synuclein give similar 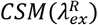 values with an average and standard deviation of & 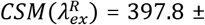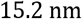 (Figures 4B and 4F). That is, we find a very similar 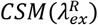 from several different single Trp proteins, differing in size, structure and physical environments (different location in peptide, ligand bound/free). This finding tracks well with our solution Trp studies. We note that the 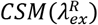 value is smaller than Trp in solution, but not outside of the calculated error. Potentially the lower 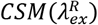 value suggests that Trp in a peptide experiences a restricted range of solvent-solute interaction energies compared to Trp in solution; *i.e.*, Trp in a peptide cannot access emissive states that are as low energy as those in solution. This is a logical conclusion given Trp in a peptide will necessarily is have restricted orientational freedom compared to bulk solvent. However, we stress the large error values on the 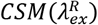 values reflecting the anticipated range of potential 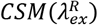 values for Trp in peptides.

These data therefore provide a ‘baseline’ range for 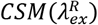, which should reflect a limiting case for the value of 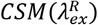 for Trp in a peptide. Fitting all our single protein Trp and solution Trp data to a shared 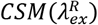 value gives 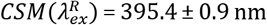. This value then represents a limiting value for 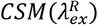 drawn from a very broad range of solvent-Trp interaction energies; it is effectively an average value. Clearly using this value of 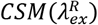 as a fixed standard for fitting Trp REES data has significant caveats. However, given the challenge of capturing meaningful data at elevated excitation wavelengths and that our modelled data (Figure S2) show Δ*G* is highly tolerant to variation in 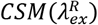, we have chosen to use this value with the much more complex data sets involving multi-Trp proteins (below). Clearly for multi-Trp proteins the extracted REES effect will be an average across all solvent-Trp environments and so the use of a well parameterised average value of 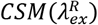 is logical. We discuss the potential for experimentally accessing 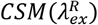 below.

### Multi-Trp proteins

Having established a limiting value of 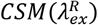 we now explore multi-Trp proteins. We have recently demonstrated that the protein REES effect can be used to predict changes in stability of multi-Trp proteins, most notably even for proteins with very large numbers of Trp residues such as monoclonal antibodies.^14^ We wish to explore whether Eq 7 retains this predictive power and to probe its sensitivity.

Figure 5 shows the temperature dependence of Δ*G* for a therapeutic mAb (IGg4-based; 150 kDa; 22 Trp residues), which is in commercial development. Figure 5A shows differential scanning calorimetry (DSC) data for the mAb, which shows *T*_m_ *onset* at 60 °C, followed by two separate unfolding transitions at 67.2 and 82.9 °C. The data shown in Figure 4B are the result of fitting the REES data to Eq 7 using 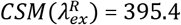 as discussed above.

**Figure 5.**
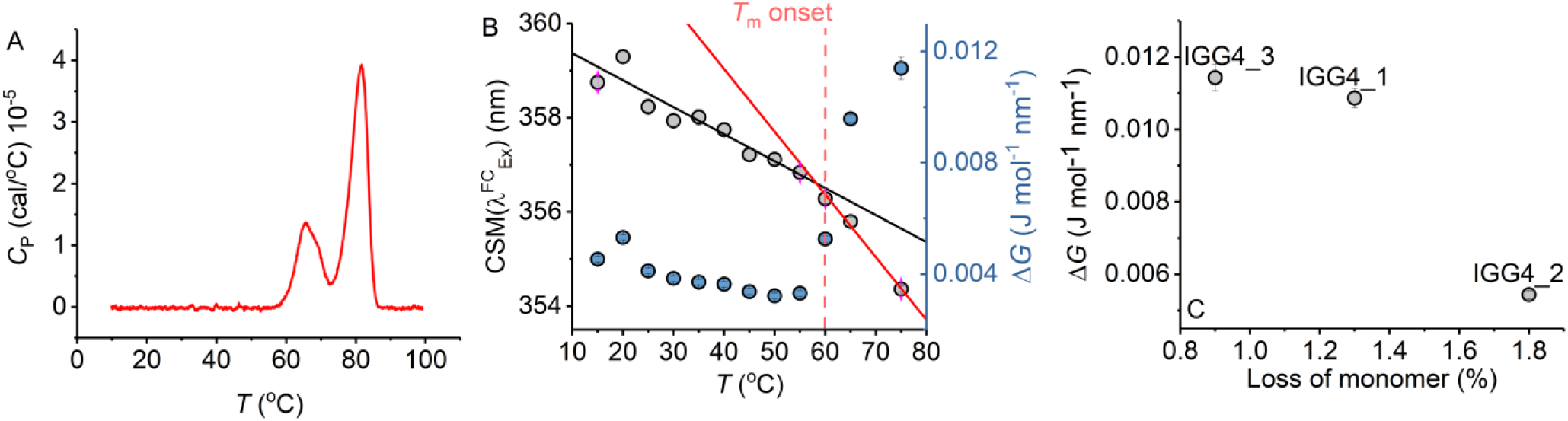
Antibody stability prediction and the effect of temperature. **A,** Differential scanning calorimetry data for mAb1. **B,** Temperature dependence of parameters extracted from fitting the IgG1 REES data to Eq 7. **C,** Percentage loss of monomer for mAb1-3 after 6 months incubation at room temperature *versus* ΔG extracted from fitting REEs data to Eq 7 at *t* = 0. Raw REES data from panel C as reported previously in ref 14.

From Figure 5B, we find that as the temperature increases Δ*G* decreases approximately linearly to ~60 °C (red dashed line) and with an approximately invariant 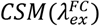 within the error of the measurement. This temperature tracks with the identified *T*_m_ *onset* from the DSC data (Figure 5A). At > 60 °C we find that 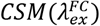 increases from 354.3 ± 0.1 at 55 °C to 359.1 ± 0.2 at 75 °C. This increase in 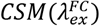 is accompanied by a larger decrease in Δ*G,* with ΔΔ*G* = 0.0042 J mol^−1^ nm^−1^ between 55 and 75 °C, compared to ΔΔ*G* = 0.0032 J mol^−1^ nm^−1^ between 15 and 55 °C. That is, we observe a breakpoint in the temperature dependence of Δ*G* (shown as the solid fitted lines). For the 15-55 °C range, we find the temperature dependence of Δ*G* is −0.1 × 10^−3^ J mol^−1^ nm-^1^ K^−1^, precisely as we found for the Trp in solution (Figure 2G). For the 55-75 °C range, this value becomes larger –-0.25 × 10^−3^ J mol^−1^ nm-^1^ K^−1^. Thus, as the protein unfolds we find an increase in 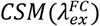 and a decrease in Δ*G*, exactly as with the chemically denatured NEMO (above). These data therefore demonstrate the sensitivity of the protein REES effect, fit using Eq 7, to altered conformational states.

Notionally, changes in the equilibrium of conformational states should track with protein stability. That is, as the free energy landscape flattens, more discrete conformational states become accessible (*i.e.*, a broader equilibrium of conformational states), including those corresponding to non-native conformations. For highly structurally similar proteins, we therefore anticipate that a decrease in the magnitude of Δ*G* will correlate with a less thermodynamically stable protein. Figure 5C shows the magnitude of Δ*G* for 3 monoclonal antibodies, in active development and all based on a common scaffold (IgG4), in relation with the fractional loss of monomer over 6 months at room temperature (reported recently^14^). From Figure 5C, we find that a decrease in the magnitude of Δ*G* correlates with a decrease in protein stability (as predicted). These data therefore suggest that not only is the magnitude of Δ*G* sensitive to the very earliest stages of protein unfolding, but also to differences in thermodynamic stability.

We have explored a similar temperature relationship with the hyperthermophilic, tetrameric, glucose dehydrogenase from *Sulfolobus solfataricus*, *ss*GDH. The natural operating temperature of the *S. Solfataricus* is ~77 °C; *ss*GDH is extremely thermally stable even at elevated temperatures and show very high rigidity relative to a comparable mesophilic protein.^29^ Figure 6A shows the far-UV circular dichroism data for *ss*GDH at a range of different temperatures. From Figure 5C there is some change in helical content with respect to temperature, most noticeable from the spectra at >85 °C. Figure 6B shows the change in ellipticity at 222 nm (*Φ*_222nm_) with respect to temperature. The solid red line in Figure 5B shows the fit to,

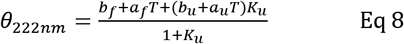

where,

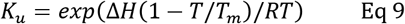

where *a* and *b* are the slope and intercept of the folded (f) and unfolded (u) baseline, respectively. *T*_m_ is the melting temperature and Δ*H* is the van’t Hoff enthalpy of unfolding at *T*_m_. From Figure 6B there is no evident complete unfolding transition and so we have restrained the parameters in Eq 8 to give a sense of where the unfolding transition would occur and an indicative *T*_m_. That is, we fix the ellipticity and gradient of the ‘unfolded’ limb of the slope to zero, which is a reasonable approximation. Fitting the data using Eq 8 gives *T*_m_ = 105.5 ± 5.5 °C. That is, the data fits to an unfolding transition that is at an experimentally inaccessible temperature. We note the significant linear slope of the ‘folded’ limb of Figure 6B. This linear phase of the thermal melt does not reflect unfolding and there is no clear consistent interpretation of the magnitude of *a*f; it is essentially always removed from analysis.^30^ Potentially it reflects changes in solvent dynamics with respect to temperature, or more trivial effects. The transition from this linear phase to the apparent unfolding transition is at ~ 80 °C.

**Figure 6.**
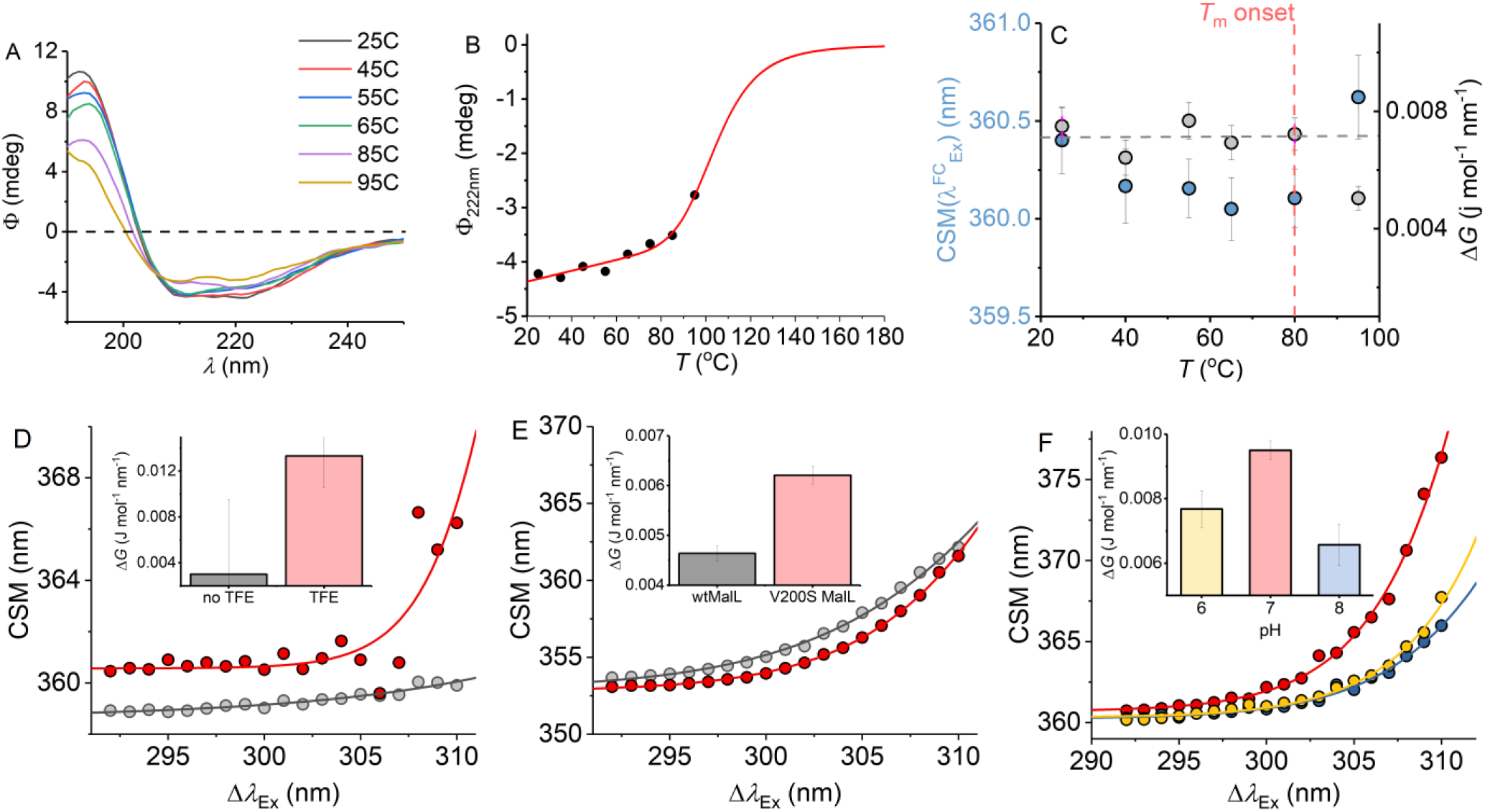
**A-C,** temperature dependence of the ssGDH REES effect and correlation with unfolding. **A,** Temperature dependence of far-UV CD spectra. **B,** Temperature dependence of Φ_222nm_. Solid line is the fit to Eq 8 as described in the main text. **C,** Temperature dependence of parameters extracted from fitting the ssGDH REES data to Eq 7. **D-F,** REES effect of C45 in the presence and absence of TFE, raw data as ref 17 (**D**), wtMalL and V200S MalL, raw data as ref 16 (**E**) and ssGDH at different pH values (**F**). The inset bar charts (**D-F**) show the magnitude of Δ*G* extract from fitting to Eq 7.

From Figure 6C we find that the magnitude of Δ*G* is essentially invariant with respect to temperature (within the error of the extracted value) up to 80 °C. As with mAb1, 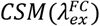 shows a small decrease with respect to temperature to 80 °C (<0.5 nm). As the notional unfolding transition occurs (95 °C), Δ*G* decreases and 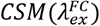 decreases. These trends are consistent with our observations with mAb1 above. However, *ss*GDH does not show the same decrease of Δ*G* with respect to temperature below the start of the unfolding transition as was evident with mAb1 and also from the anticipated temperature dependence of Δ*G* from our solution Trp studies (Figure 2G). This finding implies that whilst we anticipate the Trp REES effect will be temperature dependent, it will be protein specific. Our data do not suggest an immediate physical model for the temperature dependence of the REES effect in different proteins. However, our data potentially point to a more rigid protein (ssGDH *vs* mAb1) having a less temperature-dependent Δ*G* at temperatures below any unfolding transition. The hypothesis that more rigid protein will have a less temperature dependent REES effect seems logical given our findings of the sensitivity of the protein REES effect to even subtle changes in the equilibrium of protein conformational states. We are able to more directly explore the trend in Δ*G* on changes in molecular flexibility by correlating with evidence from NMR, simulation and pH variation. We have recently demonstrated that a *de novo* heme peroxidase (C45; four α-helix bundle; 3 Trp residues) can be rigidified (and stabilised) in the presence of 2,2,2-Trifluoroethanol (TFE).^17^ The NMR spectra (^1^H-^15^N TROSY-HSQC) show an increase in the number and sharpness of peaks in the presence of TFE, which is indicative of a more rigid protein.^17^ This rigidification also tracks with an increase in thermal stability. ^17^ Fitting the REES data to Eq 7 (shown in Figure 6D) gives a Δ*G* value that is measurably larger outside of error in the presence of TFE; Δ*G* = 0.003 ± 0.001 and 0.013 ± 0.004 J mol^−1^ nm^−1^ in the absence and presence of TFE, respectively.

For our multi-Trp examples above we are not able to rule out conformational change convolved with changes in rigidity/flexibility. Maltose-inducible α-glucosidase (MalL) has become a paradigmatic enzyme for studying the temperature dependence of enzyme activity.^31^ A single amino acid variant (V200S) is sufficient to increase the optimum temperature of reaction (*T*_opt_) from 58 °C to 75 °C, as well as having an unfolding transition at a higher temperature.^31^ Molecular dynamics simulation show that V200S is globally more rigid than the wild-type (wt) enzyme, despite the X-ray crystal structures being essentially identical.^31^ Therefore, by using MalL we are able to explore the effect of changes in protein rigidity alone on the REES effect. Fitting the extracted REES data to Eq 1 (shown in Figure 6E) gives a Δ*G* value that is measurably larger outside of error; Δ*G* = 0.006 ± 0.0002 and 0.004 ± 0.0002 J mol^−1^ nm^−1^ for V200S MalL and wtMalL, respectively.

Finally, we have explored pH variation with ssGDH. From our temperature studies (Figure 6A-C), we find that ssGDH is extremely structurally stable. In an effort to perturb the stability of ssGDH we have explored pH variation. Figure 6F shows the resulting REES data fit to Eq 7 for ssGDH incubated at pH 6, 7 and 8. From Figure 5F *inset,* we find that the magnitude of Δ*G* is largest at pH 7, with a rather lower values at pH 6 and lowest at pH 8. From our data with the mAbs, C45 and MalL, we find that a larger magnitude of Δ*G* suggests a less flexible protein. Figure S4 shows the pH dependence of the dynamic light scattering (DLS) profile. From these data we cannot identify any oligomeric change associated with pH variation. However, the DLS profiles show some variation in width, which might suggest a shift in the distribution of conformational states. These data do not obviously correlate with our REES data (Figure 5F), but potentially highlight the sensitivity of the REES data to capture changes in the equilibrium of conformational states which wouldn’t otherwise be obvious.

In summary, our combined data with multi-Trp proteins (mAbs, *ss*GDH, C45 and MalL) are consistent with the finding that a decrease in the magnitude of Δ*G* is associated with an increase in flexibility. Moreover, and as expected, reductions in molecular flexibility are correlated with increased stability. Finally, via the change in the 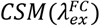 term we are able to use the fitting to Eq 7 to separately differentiate changes in molecular flexibility with unfolding. Our data therefore suggest the REES effect is potentially highly sensitive to changes in molecular flexibility outside of conformational change, as with our findings from MalL. These data therefore point to the sensitivity of monitoring the protein REES effect in multi-Trp proteins, quantified using Eq 7.

## Conclusions

The REES effect is a drastically underutilised analytical tool, given it’s potential to sensitively track changes in protein micro-states. Developing the theoretical models used to understand the effect has high potential to enable the REES effect to be used for unique applications in protein and bio-molecular analysis. For example, Kabir *et al* have recently posited a model for tracking the REES effect of a fluorescent ligand, potentially enabling the dissection of ‘hidden’ ligand bound states of proteins.^19^ Further, we have demonstrated that quantifying the REES effect allows prediction of mAb stability and this has potential for increasing the speed of drug development.^14^

Our data suggests the model presented here (Eq 7), represents a practically applicable, sensitive framework for quantifying the protein REES effect, based on fundamental thermodynamic theory. Specifically, we find that the magnitude of Δ*G* is sensitive to changes in molecular dynamics without structural change of the protein and specifically appears to be sensitive to changes in protein conformational sampling. Moreover, via the additional information pro-vided by the 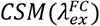 term, the model appears sensitive to early stage unfolding events and shows predictive power in assess protein stability. We anticipate Eq 7 could be modified to account for known numbers and locations of Trp residues (such as solvent accessible surface area and local protein molecular dynamics). Such data could be incorporated in Eq 7 e.g., as a weighting criterion to enable Δ*G* to be used as an independent metric of stability. Further, with the advent of a large number of high-resolution protein structures, there is very high scope for the use of homology models to fulfil this purpose where specific structures are not available.

Our model defines a maximum red shift for a given system,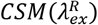, which is determined by the fluorophore and its environment. Practically, there is challenge in monitoring a low signal to noise emission spectrum at the elevated excitation wavelengths required to approach 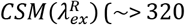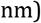, based on the range identified from our experiments. Figure 6 shows modelled power requirements to achieve an equivalent intensity emission signal. From Figure 7, the power requirement is effectively an exponential increase. That is, to accurately characterise 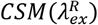 would require ~ 0.5 mW at *λ*_Ex_ = 330 nm. We note the typical output of commonly used monochromated flash lamps is ~μW. However, with the rapid development and commercial availability of high-power, stable UV LEDs, high-intensity two/three-photon laser excitation, and laser driven UV light sources we anticipate this should be practically possible.

**Figure 7.**
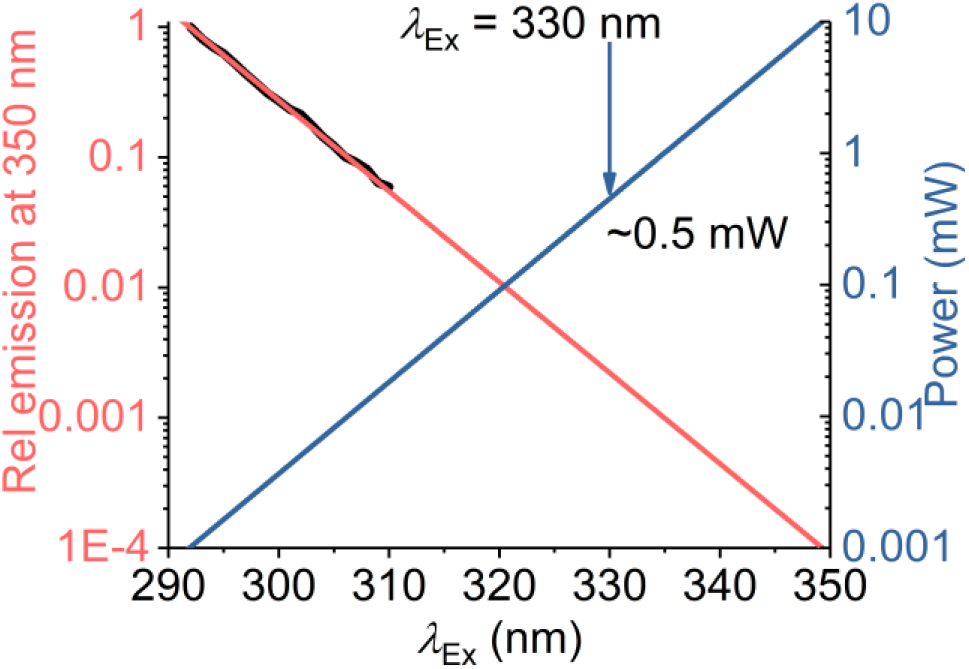
Calculated excitation power requirements to extend protein REES measurements to *λ*_Ex_ > 310 nm. The black line is the experimentally extracted (natural logarithm) excitation spec-trum of protein Trp (single Trp of NEMO as ref 15). The red line is the fit to a linear function. The blue line is the calculated power required to achieve an equivalent emission intensity at increasing values of *λ*_Ex_.

## Methods

### REES data collection

All fluorescence measurements were performed using a Perkin Elmer LS50B Luminescence Spectrometer (Perkin Elmer, Waltham, MA, USA) or an Agilent Cary Eclipse fluorescence spectrometer (Agilent, Santa Clara, CA, USA) connected to a circulating water bath for temperature regulation (1 °C). Samples were thermally equilibrated by incubation for 5 minutes at the given conditions prior to recording measurements. For all samples, the corresponding buffer control was subtracted from the spectra for each experimental condition. REES data were collected as described previously.^14^ Power readings were collected with a power meter and a UV extended silicon photodiode (Thorlabs).

### CD and DLS data collection

CD data were collected on an Applied Photophysics circular dichroism spectrometer. Corresponding buffer baselines were subtracted for each measurement. DLS data were collected on a Malvern Panalytical Zetasizer using a 50 μl quartz cuvette, thermostated to 25 °C.

### Protein preparation

α-synuclein. *ss*GDH, and mAb1 were expressed and purified as described previously in references 26, 16 and 14 respectively.

## ASSOCIATED CONTENT

### Supporting Information

Variation in dielectric and viscosity of differing methanol concentrations/temperatures, sensitivity of Eq 7 to variation in 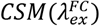, temperature dependence of 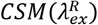, DLS data for ssGDH. This material is available free of charge via the Internet at http://pubs.acs.org.

## AUTHOR INFORMATION

### Author Contributions

The manuscript was written through contributions of all authors. All authors have given approval to the final version of the manu-script.

## ABBREVIATIONS

REES: red edge excitation shift
Trp: tryptophan
CSM: centre of spectral mass

---

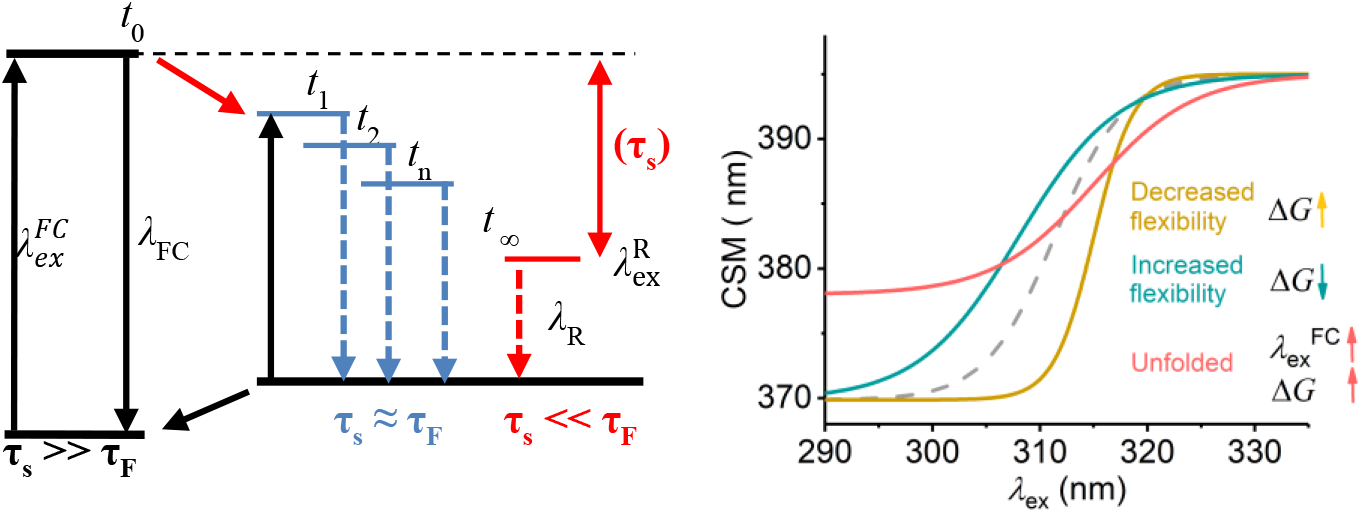

**Figure S1.**
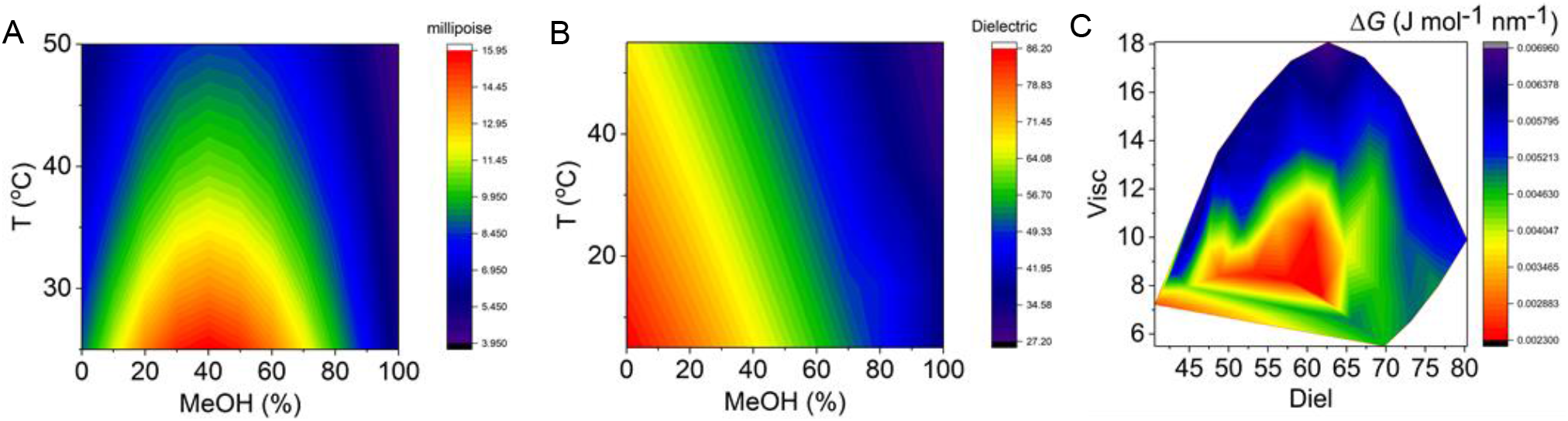
**A** and **B**, Variation in viscosity and dielectric on varying MeOH and temperature. **C**, combined viscosity and dielectric dependence of Δ*G*.

**Figure S2.**
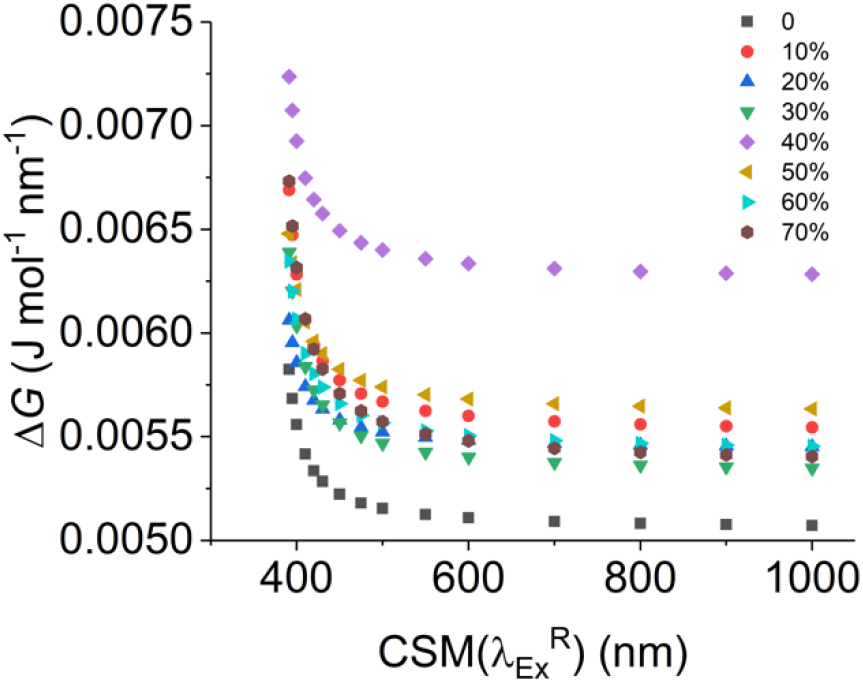
Dependence of Trp Δ*G* on 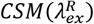 at different MeOH concentrations (20 °C).

**Figure S3.**
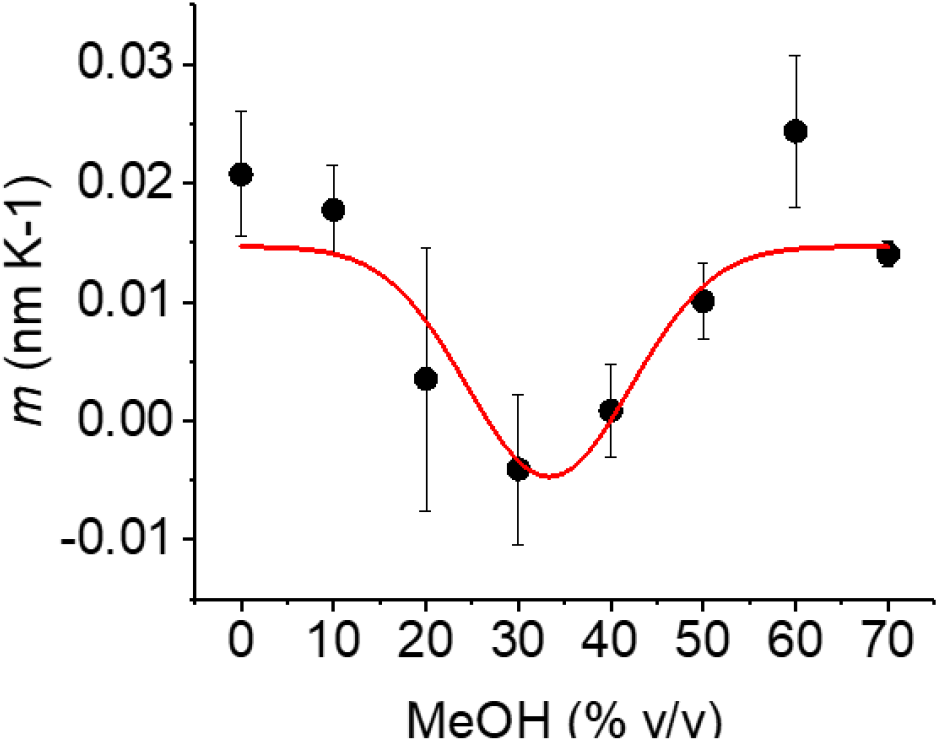
Temperature dependence of 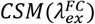 at different MeOH concentrations from Figure 2F. Where, *m* is the gradient of the fit of 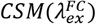 *versus T* to a simple linear function.

**Figure S4.**
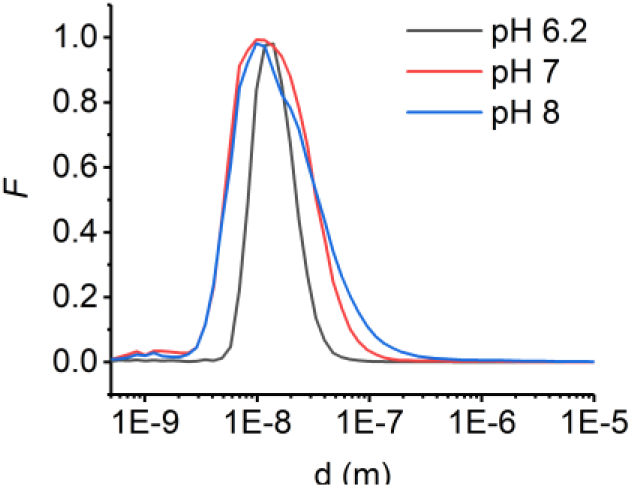
Dynamic light scattering profiles for *ss*GDH incubated at different pH values.

## REFERENCES

1. Andrey Karshikoff, Lennart Nilsson and Rudolf Ladenstein. 2015. Rigidity versus flexibility: the dilemma of understanding protein thermal stability. FEBS. J. 282, 3899–3917.

2. M Vihinen. 1987. Relationship of protein flexibility to thermostability. Protein Eng. 1, 477–480.

3. Thomas J Magliery 1, Jason J Lavinder, Brandon J Sullivan. Protein stability by number: high-throughput and statistical approaches to one of protein science’s most difficult problems. 2011. Curr. Opin. Chem. Biol. 15, 443–451.

4. Kossiakoff AA.1986. Protein dynamics investigated by neutron diffraction. Methods Enzymol. 131, 433–447.

5. Azumi T and Itoh K-i. 1973. Shift of emission band upon excitation at the long wavelength absorption edge. 1. A preliminary survey for quinine and related compounds. Chem. Phys. Lett. 22, 395–399.

6. Itoh K-i and Azumi T. 1975. Shift of emission band upon excitation at the long wavelength absorption edge. 2. Importance of the solute–solvent interaction and the solvent reorientation relaxation process. J Chem. Phys. 62, 3431.

7. Azumi T, Itoh K-i, and Shiraishi H (1976) Shift of emission band upon the excitation at the long wavelength absorption edge. III. Temperature dependence of the shift and correlation with the time dependent spectral shift. J. Chem. Phys. 65, 2550.

8. Demchenko AP. 2002. The red-edge effects: 30 years of exploration. Luminescence. 17,19–42.

9. Lippert Von E. 1957. Spektroskopische bistimmung des dipolmomentes aromatischer verbindungen im ersten angeregten singulettzustand. Z. Electrochem. 61, 962–975.

10. Mataga N, Kaifu Y, Koizumi M. 1956. Solvent effects upon fluorescence spectra and the dipole moments of excited molecules. Bull Chem Soc Jpn. 29, 465–470.

11. Chattopadhyay A, Haldar S. 2014. Dynamic insight into protein structure utilizing red edge excitation shift. Acc. Chem. Res. 47, 12–19.

12. Demchenko AP, Ladokhin AS. 1988. Red-edge-excitation fluorescence spectroscopy of indole and tryptophan. Eur. J. Biophys. J. 15, 369–379.

13. Adman ET, Jensen LH. 1981. Structural features of azurin at 2.7 Å resolution. Isr. J. Chem. 21, 8–12.

14. Knight MJ, Woolley RE, Kwok A, Parsons S, Jones HBL, Gulácsy CE, Phaal P, Kassaar O, Dawkins D, Rodriguez E, Marques A, Bowsher L, Wells SA, Watts A, van den Elsen JMH, Turner T, John O’Hara J, Pudney CR. 2020. Biochem. J. 477, 3599–3612.

15. Catici DAM, Amos HE, Yang Y, van den Elsen JMH and Pudney CR (2016) The red edge excitation shift phenomenon can be used to unmask protein structural ensembles: implications for NEMO– ubiquitin interactions. FEBS J. 283, 2272–84.

16. Jones HBL, Wells SA, Prentice EJ, Kwok A, Liang LL, Arcus VL and Pudney CR. 2017. A complete thermodynamic analysis of enzyme turnover links the free energy landscape to enzyme catalysis. FEBS J. 284, 2829–42.

17. Hindson SA, Bunzel HA, Frank B, Svistunenko DA, Williams C, van der Kamp Mw, Mulholland AJ, Pudney CR, and J. L. Anderson R. 2021. Rigidifying a De Novo Enzyme Increases Activity and Induces a Negative Activation Heat Capacity. ACS Catalysis. 11, 11532–11541.

18. Gulácsy CE, Meade R, Catici DAM, Soeller C, Pantos GD, Jones DD, Alibhai D, Jepson M, Valev VK, Mason JM, Williams RJ, Pudney CR. 2019 Excitation-Energy-Dependent Molecular Beacon Detects Early Stage Neurotoxic Aβ Aggregates in the Presence of Cortical Neurons. ACS Chem Neurosci. 10, 1240–1250.

19. Kabir, M.L.; Wang, F.; Clayton, A.H.A. 2021. Red-Edge Excitation Shift Spectroscopy (REES): Application to Hidden Bound States of Ligands in Protein–Ligand Complexes. Int. J. Mol. Sci. 22, 2582.

20. Reshetnyak YK, Koshevnik Y & Burstein EA. 2001. Decomposition of Protein Tryptophan Fluorescence Spectra into Log-Normal Components. III. Correlation between Fluorescence and Microenvironment Parameters of Individual Tryptophan Residues. Biophys. J. 81,1735–58.

21. Hammond, G. S. 1955. A Correlation of Reaction Rates. J. Am. Chem. Soc. 77, 334–338.

22. Barczewski AH, Ragusa MJ, Mierke DF, Pellegini M. 2019. The IKK-binding domain of NEMO is an irregular coiled coil with a dynamic binding interface. Sci Rep. 9, 2950.

23. Meade, R.M., Fairlie, D.P. & Mason, J.M. 2019. Alpha-synuclein structure and Parkinson’s disease – lessons and emerging principles. Mol Neurodegeneration 14, 29.

24. Tuttle MD, Comellas G, Nieuwkoop AJ, Covell DJ, Berthold DA, Kloepper KD, Courtney JM, Kim JK, Barclay AM, Kendall A, Wan W, Stubbs G, Schwieters CD, Lee VM, George JM, Rienstra CM. 2016. Solid-state NMR structure of a pathogenic fibril of full-length human α-synuclein. Nat. Struct. Mol. Biol. 23, 409–415.

25. Jain N, Bhasne K, Hemaswasthi M, Mukhopadhyay S. 2013. Structural and Dynamical Insights into the Membrane-Bound α-Synuclein. PLOS One. 8, e83752.

26. Meade RM, Morris KJ, Watt KJC, Williams RJ, Mason JM, 2020. The Library Derived 4554W Peptide Inhibits Primary Nucleation of α-Synuclein. J. Mol. Biol. 24, 166706.

27. Catici DAM, Horne J E, Cooper GE & Pudney CR (2015) Polyubiquitin drives the molecular interactions of NF-κB essential modulator by allosteric regulation. J. Biol. Chem. 290, 14130–9.

28. Fenner BJ., Scannell M, and Prehn JHM. 2010. Expanding the substantial interactome of NEMO using protein microarrays. PLoS One. 5, e8799.

29. Vieille C, Zeikus GJ. 2001. Hyperthermophilic enzymes: sources, uses, and molecular mechanisms for thermostability. Microbiol. Mol. Biol. Rev. 65, 1–43.

30. Greenfield NJ. 2006. Using circular dichroism collected as a function of temperature to determine the thermodynamics of protein unfolding and binding interactions. Nat Protoc. 1, 2527–35.

31. Hobbs JK, Jiao W, Easter AD, Parker EJ, Schipper LA, Arcus VL. 2013. Change in Heat Capacity for Enzyme Catalysis Determines Temperature Dependence of Enzyme Catalyzed Rates. ACS Chemical Biology. 8, 2388–2393.

